# Molecular Dynamics of Phosphatidylcholine Model Membranes of Splenic Lymphoma Cells

**DOI:** 10.1101/2024.07.23.604722

**Authors:** Michael Kio, Joshua Lucker, Jeffery B. Klauda

**Affiliations:** Department of Chemical and Biomolecular Engineering, University of Maryland, College Park, MD 20742, USA; Institute for Physical Science and Technology, Biophysics Program, University of Maryland, College Park, MD 20742, USA

## Abstract

In eukaryotic cell membranes, phosphatidylcholine (PC) is one of the most prevalent phospholipids. Using the CHARMM36 lipid force field, we conducted molecular dynamics (MD) simulations on phosphatidylcholine (PC) only headgroup with varied fatty acid chains. Specifically, we investigated five PC components: 1,2-diauroyl-sn-glycero-3-phospocholine (DLPC), 1,2-dimyristoyl-sn-glycero-3-phosphocholine (DMPC), 1,2-dipalmitoyl-sn-glycero-3- phospcholine (DPPC), 1-palmitoyl-2-oleoyl-sn-glycero-3-phosphocholine (POPC), and 1- stearoyl-2-oleoylphosphatidylcholine (SOPC) in both pure and cancer model of PCs. We analyzed various characteristics such as lipid area, lateral compressibility, deuterium order parameter, bilayer thickness, radial distribution functions, and electron density. Our research revealed that PCs in the cancer model membrane are larger in surface area per lipid compared with pure PC membranes. This suggests that our PC model for cancer PCs may be more permeable and porous compared to pure PC membraness. In general, chain order parameter values were lower in cancer PCs compared to pure PCs. The electron density studies of cancer PCs revealed a decrease in bilayer thickness as temperature increases, indicating that cancer PCs experience thinning at higher temperatures. Overall, our results give insight into significant differences between the cellular makeup and functioning of pure PCs and cancer PCs at the molecular level.

## 1 Introduction

Biological membranes consist of lipids and proteins that form a lipid bilayer structure and are essential to cells^1^. These can enclose a cells, separates the inside and outside of a cell, and selectively permits or restricts the passage of molecules. This thin layer contains transmembrane proteins that provide additional function to the cell membrane. Therefore, knowledge about the properties of lipid bilayers is imperative to comprehend biology better and is useful for various medical and pharmaceutical applications such as biosensors^2, 3^ and drug delivery systems specifically, ATP-binding cassette (ABC) transporters^4–6^. However, we still lack detailed information regarding its microscopic elements and behaviors. Experimentally measuring the thickness within the bilayer (which is only ∼4 nm thick) is extremely hard because of extreme variability, disorderliness, and mobility. Molecular dynamics (MD) simulations have been beneficial in analyzing certain aspects of this membrane at a nanoscopic level ^7–9^. Lipids are a vital component of cellular membranes, their amphiphilic structure forming a barrier between the cell and any external environment^10^. They can also facilitate transport processes via proteins by forming channels, pumps, and gates ^11^. Although the components that make up these membranes may vary depending on the type of cell, studying lipid diversity can give useful insight^12^. In the biologically significant disordered phase, membrane structures remain dynamic, as the lipids constantly change conformation, diffuse in a lateral plane, rotate, and move collectively^13^. This movement spans various time scales from ps to ms^14^.

A wide range of chemically distint lipids constitute the mammalian plasma membranes ^15^. These lipids can be roughly divided into three groups. The first group consists of glycerophospholipids. When broken down by their head groups, they include phosphatidylcholine (PC), which is the study focus, phosphatidyl-myo-inositol (PI), phosphatidylserine (PS), and phosphatidylethanolamine (PE). Each headgroup has variable tail groups; this includes the number and locations of double bonds and length of their acyl chains. The structure of the phospholipid bilayers is greatly affected by these distinctions. The second group is cholesterol, while the third is composed of sphingolipids, represented mostly by sphingomyelin (SM). Cholesterol aids in forming liquid ordered phase by segregating saturated tails and SM lipids. This effect is universal among phospholipid-cholesterol combinations, no matter the amount of unsaturation in the sn–2 tail^16–18^. The makeup of plasma membranes varies a lot from one type of cell to another. Cells are believed to acquire necessary properties for their membranes by controlling the composition of lipids. Moreover, there have been noticeable discrepancies between lipid headgroups in pure and cancer cells since 1960s ^19–25^. Following that, concentrations of glycerophospholipids, sphingomyelin, and cholesterol have been calculated individually in rat^23, 24^, mouse ^22, 26^, and human cells ^21, 22^.

Our interest in this work is to focus on investigating the properties of cancer membrane models. The plasma membrane lipid concentrations in healthy and cancer PCs can vary greatly based on cell species. However, individual cell species tend to have the same trends in changes independent of mammalian differences. There has been an interesting observation regarding changes in lipid concentrations found in these membranes ^20–31^. Researchers report that the level of cholesterol relative to total phospholipids (C/PL ratio) was 0.74 (42.5 mol%) in healthy cells and as low as 0.32 (24.2 mol%) in leukemic membranes ^24, 26^. This decrease in cholesterol concentration is also seen in other species of rats and mice. In addition, head groups of component phospholipids like PE, PC, PI and PS, (lyso-PE and lyso-PC) lysophospholipids, and (SM) sphingolipids were observed with an increase in PE and a reduction in SM in leukemic mouse cells. Furthermore, measurements on the length of alkyl tail chains and number of unsaturated bonds showed a reduction in cancer cells of saturated tail chain levels from (59.3- 44.0) mol% ^26^.

This study set out to explore, at a molecular level, the physical changes in pure and cancer PCs implementing CHARMM36^32–47^ lipid force fields and CHARMM-GUI *Membrane Builder*^33^. Previous research has focused on the measurement of fluorescence depolarization in cancer and normal cell membranes^48–49^. The authors found a drop in an orientational order parameter ^28–29^ for the leukemic plasma membranes, indicating that the probe molecules were more disordered in cancer cell membranes than in normal ones. Advancements in supercomputers have made it possible to perform ns-μs MD simulations for complex and large systems like model plasma membranes. These calculations can provide data about membrane dynamics and structure, such as self-diffusion constant^51^ and lipid lateral distribution functions^52^. A MD study of PC model membranes can offer valuable information regarding their structural and dynamic characteristics important to the understanding of their role in real biological membranes. Modelling plasma membranes are complex but we will focus on simple models and representation of the PC lipids. This study examines the properties of a simplified splenic cancer PC membrane model and compares it to pure PC lipid bilayers, using atomistic MD simulations. To create the splenic PC model, we simplified the experimental data and ensure that it sufficiently describes the PC lipids in the system for accurate results^26^. Using the mole fractions obtained from lipidomics data for lymphoma PC ^25,48^. We calculated the number of lipids using the mole fraction present in pure and cancer PCs. However, due to the small mole fraction of cancer PCs observed in the experimental data, we decided to combine all cancer PC groups in our model to represent large enough PCs for meaningful result. The variables investigated include structural quantities like lipid area, lateral compressibility, deuterium order parameter, bilayer thickness, radial distribution function and electron density.

## 2 Methods

Using experimental data, simulation model membranes are created and compared focusing on pure membrane models with PC lipids in cancer (Table 1) and a model to represent the cancer PC membrane^25,48^ (Table 2) The detailed calculations and representations are shown in S.0.These pure PC models are compared the cancer PC model to investigate biophysical property changes. These models includes five PC lipids (DLPC, DMPC, DPPC, POPC, SOPC). Despite the limited amount of lipids, our approach was to augment the number of lipids within the cancer (PCs) by modelling them collectively and doing analysis for each PC component.

**Table 1.** Pure PCs simulated,, temperature, and total simulation length. All systems consisted of 72 total lipids (36 per leaflet).

| Lipid | T (°C) | T(ns) |
| --- | --- | --- |
| DMPC | 30,50,60 | 70 |
| DLPC | 20-60 | 180 |
| DPPC | 30,50,60 | 150 |
| POPC | 20-60 | 200 |
| SOPC | 20-60 | 100 |

**Table 2.**
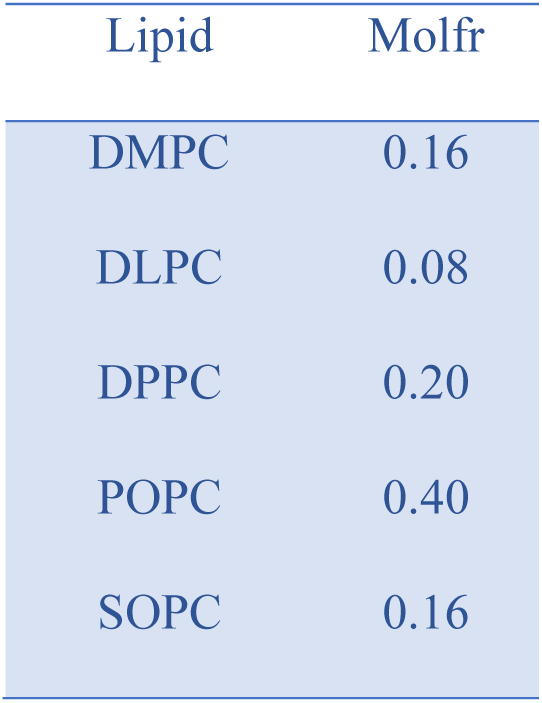
Mole fraction of lipids in the mixed PC cancer model. This system was run for 100ns and at temperatures 20-60°C in 10°C increments.

**Table 3.** The SA/Lip and KA values of experiments^27^ for single PC model and mixed cancer PC models.

| Temp | Pure POPC SA/Lip ($\text{\AA}^2$) | Mixed Cancer SA/Lip ($\text{\AA}^2$) | Exp Models of POPC<sup>27</sup> SA/Lip ($\text{\AA}^2$) | $K_A$ Pure POPC (N/m) | $K_A$ Cancer (N/m) |
| --- | --- | --- | --- | --- | --- |
| 20°C | 63.82 +/- 0.3 | 64.72 +/- 0.5 | 62.70 +/- 0.18 | 0.22 | 0.13 |
| 30°C | 64.02 +/- 0.3 | 64.98 +/- 0.5 | 64.30 +/- 0.19 | 0.19 | 0.10 |
| 40°C | 66.36 +/- 0.2 | 67.31 +/- 0.6 |  | 0.27 | 0.11 |
| 50°C | 67.84 +/- 0.2 | 68.26 +/- 0.3 | 67.30 +/- 0.21 | 0.27 | 0.14 |
| 60°C | 68.83 +/- 0.3 | 69.32 +/- 0.3 | 68.10 +/- 0.19 | 0.12 | 0.24 |

### 2.1 Simulation Details and Calculation

The CHARMM36 lipid force field (C36FF)^32^ and CHARMM-GUI Membrane Builder^33^ are implemented in the development the membrane models. Average of 40 water molecules for pure and 43 water molecules for cancer per lipid was implemented to ensure that the lipids were fully hydrated Table 1. The TIP3P water model^53–60^ with C36FF was employed, along with van der Waals interactions fading between 10 and 12Å through a force-based switching function^61^. All bond lengths involving hydrogen atoms were constrained using the SHAKE algorithm with an accuracy of 10^-5^, to constrain any bonds involving hydrogen atoms as well as the angle of the hydrogen-oxygen-hydrogen bond in water^62^, while Particle Mesh Ewald^63^ was employed for electrostatic calculations, with an interpolation order of 6 and a direct space tolerance of 10^-6^. MD simulations at varied temperatures were conducted by means of NAMD^64^; these temperatures exceeded the gel-transition points of the respective lipids. After following the standard CHARMM-GUI six minimization and equilibration steps, we ran the simulations for 70-200 ns for the various PC lipids Table 1 and 2. Both pressure and temperature were set to 1 bar and kept constant respectively, non-hydrogen atoms were regulated using Langevin dynamics, which kept the temperature constant with a coupling coefficient of 1 ps^-1^. In addition, Nosé-Hoover Langevin-piston algorithm was applied to ensure a steady pressure using a piston cycle of 50-fs and a 25-fs piston decay^65–66^. VMD ^67^ was used for snapshots of the bilayers, while Gnuplot^68^ was utilized to create structure property plots.

### 2.2 Bilayer Structural Properties

#### 2.2.1 Area of Lipid

To calculate the area per lipid, we followed previously reported studies^39^. The average area per lipid is calculated by dividing the total area of the system in the horizontal plane (xy-plane) by the amount of lipids found in each monolayer. The statistical errors of SA/lip were determined by calculating the block averages. The area compressibility K_A_ was calculated by measuring the variation of the overall surface area within the x-y plane.

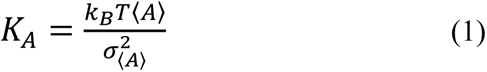

where k_B_ is the Boltzmann’s constant, 〈*A*〉 is the average total area per leaflet, *σ*^2^_〈*A*〉_ is the mean square fluctuation of that area and T is the temperature (in Kelvin)^39^.

#### 2.2.2 Multicomponent Surface Area of Lipid

The overall surface area per lipid (SA/lip) was determined by dividing the area of the simulation box by the number of lipids per leaflet. A Voronoi diagram was then generated for each system using Quickhull^69^, with each polygon representing a representative atom. Quickhull was used to calculate the component SA/lip, which involved obtaining the X and Y coordinates of representative atoms for each lipid. The component SA/lip was obtained by averaging the sum of the areas of all representative atoms for each type of lipid present in the system. Once the overall SA/lip was calculated, the KA was then determined using this formula:

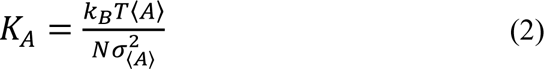

where k_B_ is the Boltzmann’s constant, 〈*A*〉 is the average total area per leaflet, N is the, the number of lipids per leaflet *σ*^2^_〈*A*〉_ is the mean square fluctuation of that area and T is the temperature (in Kelvin)^39^

#### 2.2.3 Deuterium order parameter S_CD_

The S_CD_ (specific chain orientation distribution) of each carbon links up with the average ensemble configuration and is understood to measure the overall order of the lipid bilayer, meaning a higher figure denotes making it more structured. The S_CD_ can be calculated by taking the average of the second-degree Legendre polynomial^50^. Equation is expressed as;

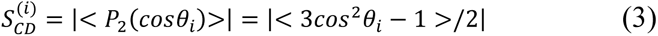

For each carbon atom connected to a hydrogen, θi represents the angle between the C-H bond and the normal of the bilayer. The angular bracket expresses the time and ensemble average. As temperatures drop, the S_CD_ of almost all carbons go up, which correlates with a thicker bilayer as well as an increased order in the chain.

#### 2.2.4 Electron Density

Electron density profile (EDP) refers to the likelihood of an electron being found at a particular position relative to the membrane cener. The EDP is determined by calculating and averaging the electron density over a certain position range. Given that our lipid bilayer systems are centered and symmetrical at z=0, an additional averaging process is used for EDP calculations in both positive and negative z directions. The z=0 mark holds the terminal methyl groups of hydrocarbon chains which has the most reduced total electron density^70–75^. The electron density profile of a typical membrane shows a dip in the center where the CH3 groups are located, two peaks at the head groups (phosphate groups), and a decrease in density at the water layer.

#### 2.2.5 2-Dimensional Radial Distribution Function (2D RDF)

Ordering of the lateral organization in a lipid bilayer can be examined using the two-dimensional radial distribution function (2D RDF). The 2D RDF is a way to analyze the distribution of atoms or molecules around a central mass and is also useful in determining the number of solvation layers present in a lipid bilayer. One method for finding this is by calculating the RDFs by identifying the nearest lipid head group atom (phosphate oxygen) and then grouping them into bins based on their distance. Additionally, RDFs can provide evidence of hydrogen bonding if the first peak appears within a specific range indicating an H-bond between the water molecules and phosphate oxygen in the headgroup.

## 3. Results and Discussion

In the following section, we will mention the findings from our MD Simulation comparing pure and cancer PCs using a 5-lipid model system. Our results will cover various aspects such as surface area of lipid, lateral compressibility, deuterium order parameter, bilayer thickness, radial distribution functions, and electron density.

### 3.1 Surface Area per Lipid and Area Compressibility Modulus

The surface area per lipid (SA/lip) and area compressibility were studied and compared between pure and cancer PC models, as well as to experimental values of oveall POPC membrane model. First, we focused on equilibration of our systems to validate the amount of time to collect data. The system are equilibrated based on SA/lip vs time plot. Fig S.1 and Fig S.2 displays the SA/lip vs time plot for pure DLPC at 30°C - 60°C and mixed cancer DLPC at 30°C – 60°C using the same force field.

Table 2 presents the experimental values^27^ and simulations for the average SA/lipid and K_A_ for Pure or Single POPC and cancer mixed membrane at 30°C-60°C and Table S.2 displays the SA/lipid and K_A_ for experimental values, pure and cancer models for the the other 4 PC components. The data on the tables indicates that cancer PCs have a significantly higher suface area per lipid than pure PCs at the same temperature and as temperature increases. This is a significant alteration, most likely caused by changes in the lipid makeup of the PCs^65^.

Table 2 also shows the values for K_A_ with results demonstrating that the cancer PCs has lower K_A_ values than the pure PCs at temperatures ranging from 20°C-60°C. However, at higher temperatures of 60°C, the K_A_ values for the cancer PCs significantly increase and surpass those of the pure PCs. The K_A_ values for the cancer PC models are considerably lower than the pure PC models, yet both are significantly lower compared to PCs in a glycerophospholipid membrane bilayer^70^.

As the temperature rises, the surface area per lipid also increases in both the pure and cancer PCs. This trend is also consistent with the findings for unsaturated POPC shown in Table 2. Additionally, the cancer PCs have a higher SA/lip ratio than the pure PCs, which aligns with previous studies on PC lipids^55^,

### 3.2 Deuterium order parameter (S_CD_)

Figure 1 displays the SCD values for different *sn*-1 and *sn*-2 parameters for both pure and cancer DMPC models at temperatures ranging from 30 to 50°C. These graphs clearly show a decrease in SCD values as the carbon number increases. When the temperature rises, the C4-C14 side chains become less ordered and more liquid-like, leading to a decreasing average SCD for all lipids. The graph shows that the average SCD of pure DMPC has the highest peak when the temperature is 30°C and at its lowest point when it reaches 60°C for pure PCs *sn*-1 and *sn*-2. This pattern is also observed in cancer PCs for tails at various temperatures (as shown in Fig S.2 and Table S.2-S.5), indicating the effect of temperature on the deuterium order parameters for both the cancer and pure PCs. These findings also indicate that the order parameter values are lower in cancer PCs when compared to pure PCs.

**Fig.1.**
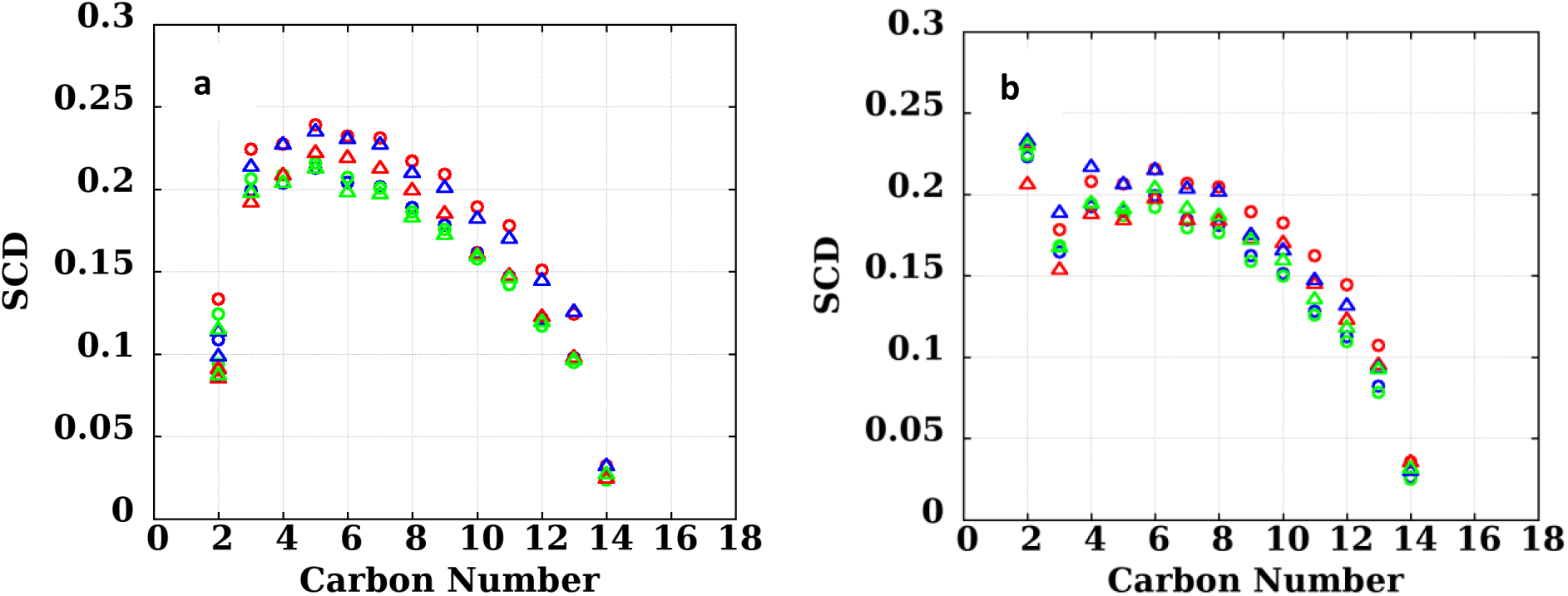
The simulated SCD of DMPC (pure-Red Circle 30°C, pure-Blue Circle 50°C, pure-Green Circle 60°C and Cancer (cancer-Red Triangle 30°C, cancer-Blue Triangle 60, cancer-Green Triangle 60°C. a) *sn*-2 b) *sn*-2.

Figure 1 depicts the simulated 5-lipid system SCD values of *sn*-1 and *sn*-2 for pure and cancer PCs at a temperature of 60°C. The results show that pure DPPC has the highest S_CD_ while DLPC has the lowest for *sn*-1. On the other hand, SOPC and DPPC have the highest S_CD_ while DLPC has the lowest for *sn*-2. In contrast for cancer PCs, DMPC and DPPC exhibit the highest S_CD_ for *sn*-1 while POPC and SOPC have the lowest. For *sn*-2, it is observed that DPPC and DMPC have the highest S_CD_ while POPC and SOPC have the lowest. This variation can be attributed to differences in carbon atom alignments between pure and malignant PCs. Despite these differences, the simulated S_CD_ values for pure PCs closely resemble experimental data ^63^. It should be noted that the cancer PCs showed a lower value of S_CD_ than pure PCs and this value decreased as unsaturation increased^68^.

Fig.2 The simulated SCD of five lipid PC system at 60°C (pure DLPC-Red Circle, pure DMPC-Blue Circle, pure DPPC-Green Circle, pure POPC-Yellow Circle, pure SOPC-Purple Circle, (cancer DLPC-Red triangle, cancer DMPC-Blue triangle, cancer DPPC-Green triangle, cancer POPC-Yellow triangle, cancer SOPC-Purple triangle. a) Pure *sn*-1, b) Pure *sn*-2, c) Cancer *sn*-1, d) *sn*-2

**Fig.2 The.**
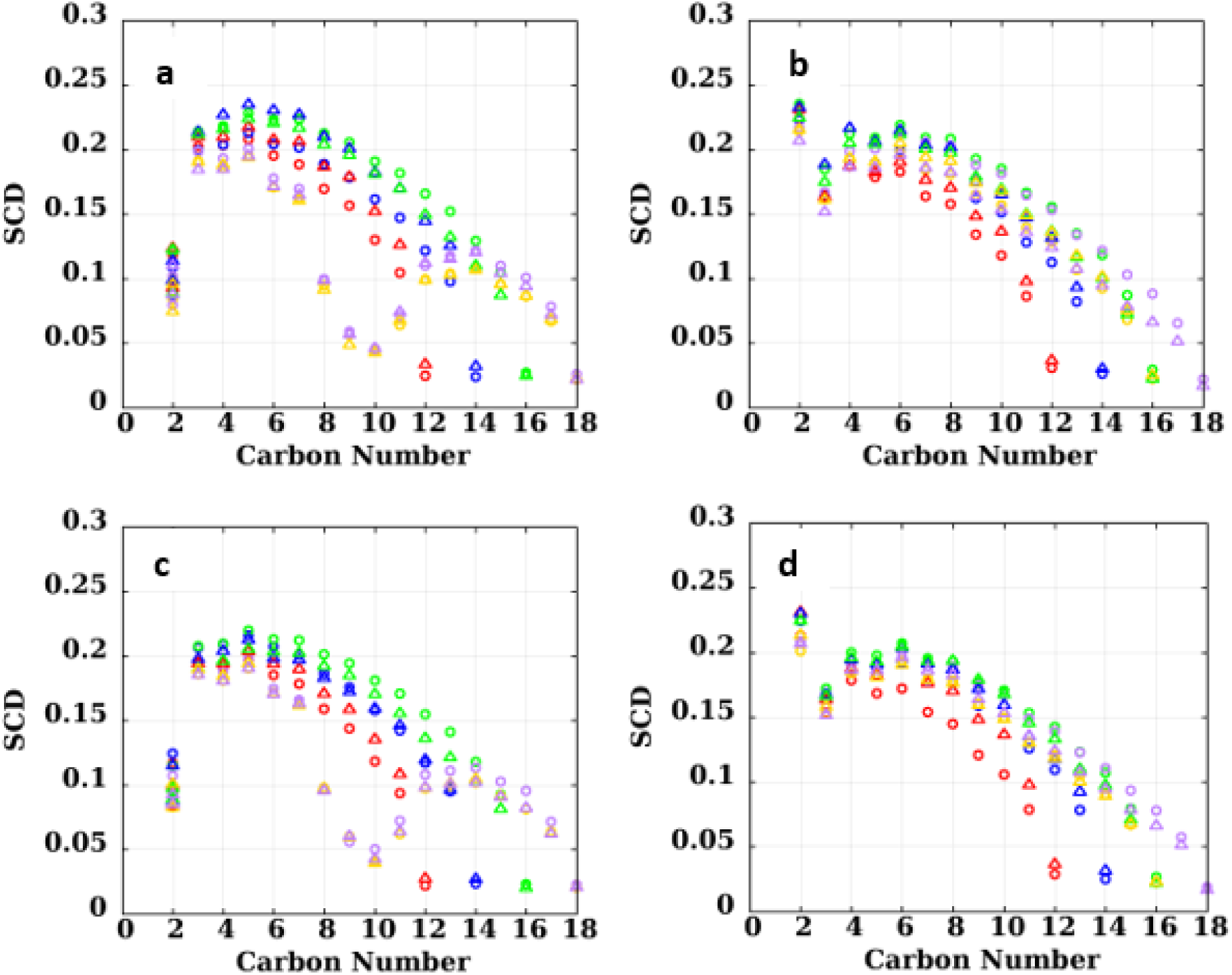
simulated SCD of five lipid PC system a) Pure *Sn-1* PC b) Pure *Sn-2* PC c) Cancer *Sn-1* PC d) Cancer *Sn-2* PC at 60°C.

In Figure 3, the slight change in S_CD_ within the 5-lipid system of pure and cancer Sn-1 and Sn-2 PC is displayed as temperature increases. The Sn-1 DMPC PC has the highest SCD at 20°C, but then drops below DPPC and DLPC at 30°C before rising again. Meanwhile DLPC, DPPC, POPC, and SOPC-all show a gradual decrease in SCD with increasing temperature. All Sn-2 lipid maintains a relatively close SCD, with a slight decrease of SCD as temperature increases from 20°C-60°C. The SCD of Sn-1 unsaturated POPC and SOPC cancer PC is significantly lower than the SCD of Sn-2, POPC and SOPC cancer PC. These findings suggest that SCD is highest at lower temperatures and decreases as temperature increases within the 5-lipid PC system. Furthermore, a higher level of unsaturation correlates with a lower order parameter and less organized membranes.

**Fig. 3.**
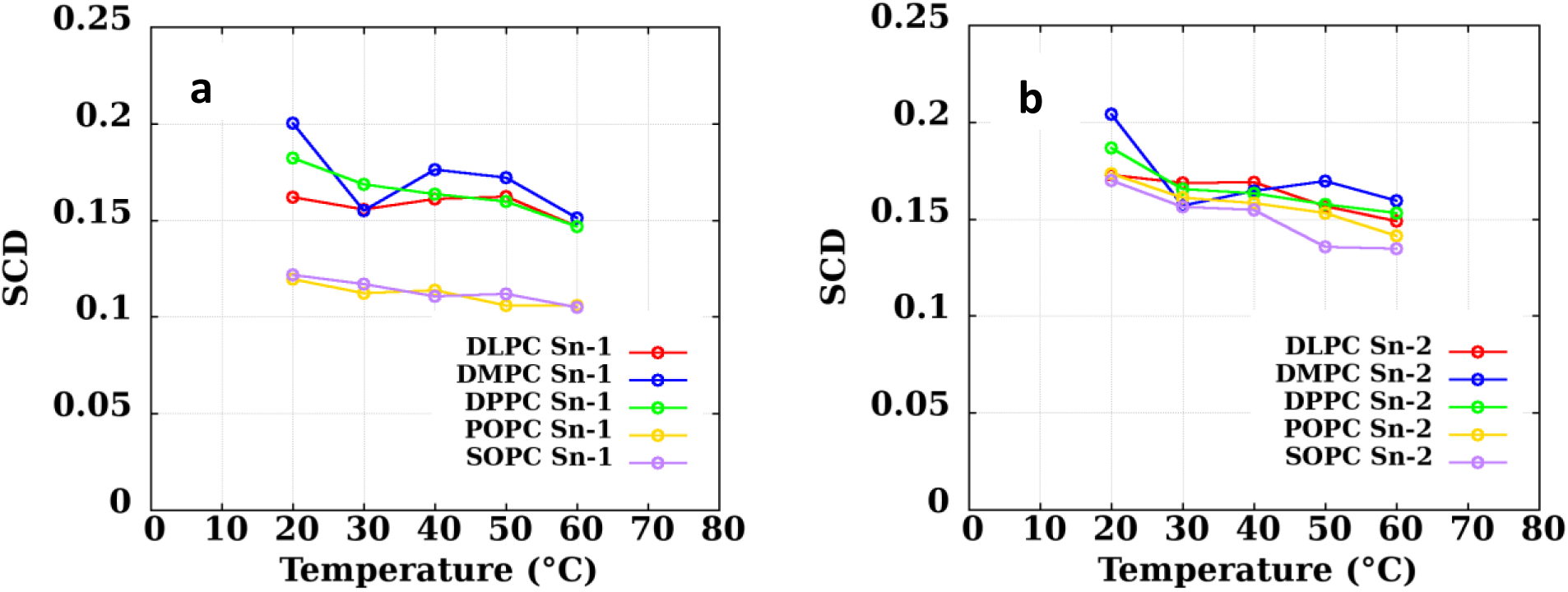
The Simulated temperature for 5-lipid system a) Sn*-1 and* b) *Sn-2* of Cancer PCs at 20°C -60°C.

The SCD values in cancer PCs were mostly lower than those of pure PCs. Moreover, the degree of order decreased as the temperature rose. The decrease in the amount of component lipids, combined with an increase in pure and cancer PC temperatures caused a noteworthy decline in the overall conformation order of the membrane. These findings suggest that the order parameter of cancer PCs were noticeably lower than that of pure PCs. This indicates a significant decrease in the conformational order of cancer PCs compared to pure ones. As the temperature increases, the degree of order decreases for SCD in almost all carbon atoms. This leads to a decrease in the overall organization of lipids in the bilayer as temperature rises, which is supported by less organized PCs.

### 3.3 Electron Density

Fig 4 shows electron density profile for DLPC for pure PC at temperatures 20°C, 30°C, 50°C 60°C. the black line (total) depicts the peaks of the pure DLPC.

**Fig 4.**
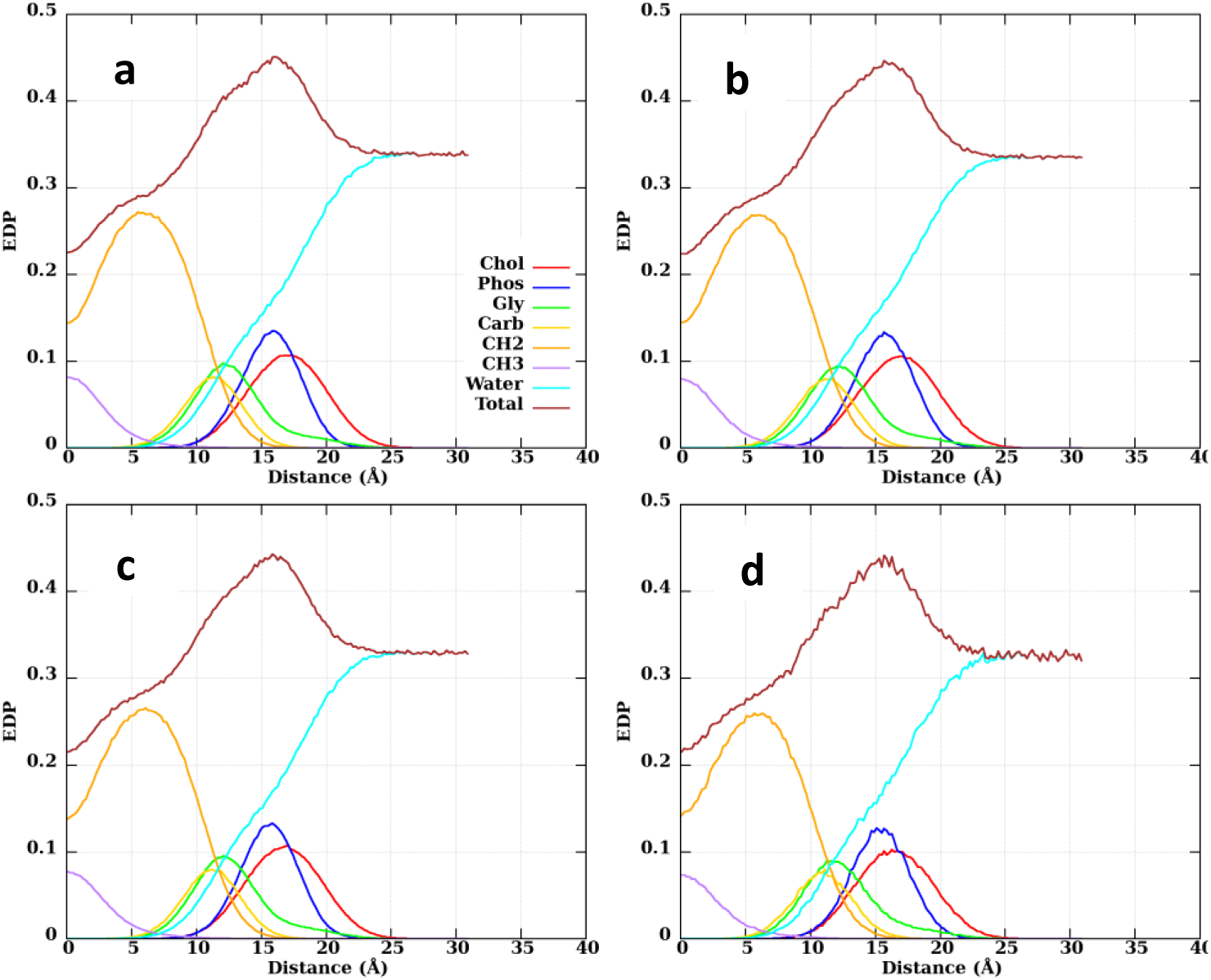
Electron density profiles for DLPC pure PC at temperatures a) 20°C b) 30°C c) 50°C d) 60°C.

At a lower temperature of 20°C, the peak is at 18A, while at 30C it shifts to 17A. As the temperature increases, the peak for 50°C moves to 16A and for 60°C, it drops even further to 15A.

This suggests that as the temperature increases, the peak distance decreases, which implies that the PC becomes thinner, confirming previous studies on DLPC thickness as shown in Fig 6. By analyzing the contributions of different moieties in the system, it can be seen that CH2 molecules and water have the highest EDP peaks, while carbon and CH3 has the lowest peak.

Figure 5 displays the electron density profile (EDP) of a 5-lipid PC system: DLPC (red line), DMPC (orange line), DPPC (purple line), POPC (yellow line), and SOPC (green line) at 40°C and 60°C. The black line (total) depicts the peaks of the cancer PC at 40°C and 60°C with the highest total electron density between 19-23Å.

The results show that at temperatures of 40°C and 60°C, Fig.3 and 50°C, Fig S.8. DMPC has the smallest EDP, remaining relatively constant between 5A and 22A before tapering off at 25A.

**Fig 5.**
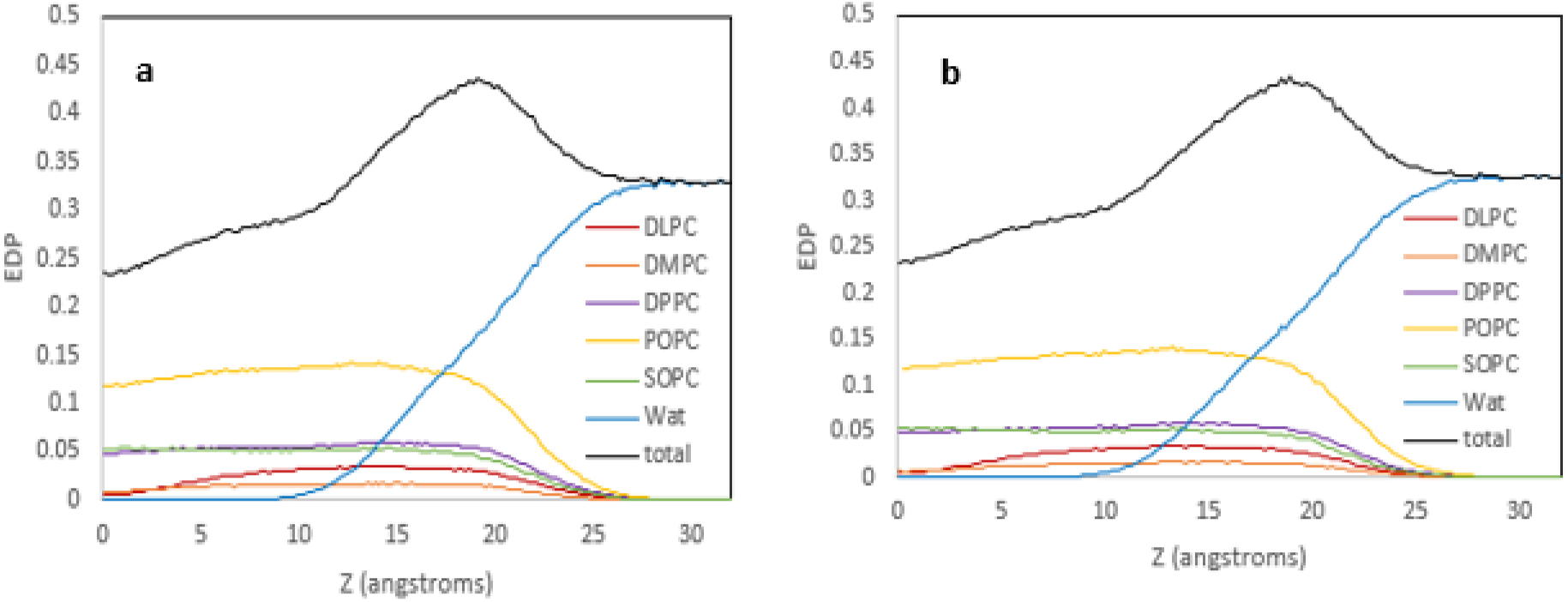
Electron density profiles for 5-lipid system of cancer PCs for temperatures a) 40°C and b) 60°C.

This is followed by DLPC, SOPC, and DPPC with similar patterns and also tapering off at around 25A. POPC has the highest EDP compared to other PCs at temperatures of 40°C and 60°C. Water is shown to have the largest contribution to the EDP in this system, with a higher EDP at 40°C compared to 60°C. As temperature increases from 40°C to 60°C Fig S.8 showing 50°C, the EDP of the 5-lipid PC system also increases. The peak of the total EDP for the five-lipid PC system is 19A at 40°C and 18A at 60°C, indicating that temperature increase results in thinning of the cancer PCs. Additionally, water exhibits a higher EDP at lower temperature 40°C than at higher temperature 60°C and the water peak in the PC system is found to be higher at 40°C compared to 60°C. This observation also validates the results from the bilayer thickness. compared to 60°C.

### 3.4 Bilayer Thickness

The bilayer thickness (D_B_) and hydrophobic thickness (2D_C_) are compared against data estimated by experiments. The simulation results agree with the experiments and the overall thickness of the bilayer PCs, showing that D_B_ and 2D_C_ reduces as the temperature increases Table S.6.

Fig. 6 makes comparisons of pure and cancer DB and 2DC for PCs of saturated DLPC, DMPC unsaturated POPC, and SOPC at temperatures ranging from 20°C-60°C. In Fig S.3 a similar comparison was made for pure and cancer saturated DPPC at temperatures ranging from 50°C-60°C. It is observed that pure and cancer unsaturated POPC and SOPC have a greater DB and 2DC than pure and cancer saturated DLPC and DMPC owing to double bonds resulting in permanent kinks in the hydrocarbon chains which take up more space. In regards to DMPC and DPPC, pure DBsim, and DCsim almost coincide with DBexp and 2DCexp; experimental values at all temperatures except in the pure DBsim of DMPC at higher temperature of 60°C^73,74^. For DLPC and POPC, pure DBsim and pure 2DC has offset with experimental values and for POPC and DMPC pure 2DC are better aligned with experimental values, with pure DMPC DBsim almost in agreement with experimental values and POPC DBsim in offset at 20°C-60°C. Cancer DB and 2DC is shown to be higher in all the 5-lipid bilayer system when compared to Pure DB and 2DC, Except for DPPC, where pure DB and 2DC is greater than cancer DB and 2DC. DBsim and 2DCsim of both saturated and unsaturated pure and cancer PCs generally decreases as temperature increases, with fluctuations observed in cancer DLPC, POPC, SOPC and pure SOPC as temperature increases from 20°C-40°C. Data of DHH is not an area of focus due to lack of available experimental DHH value.

**Fig.6 Similarities.**
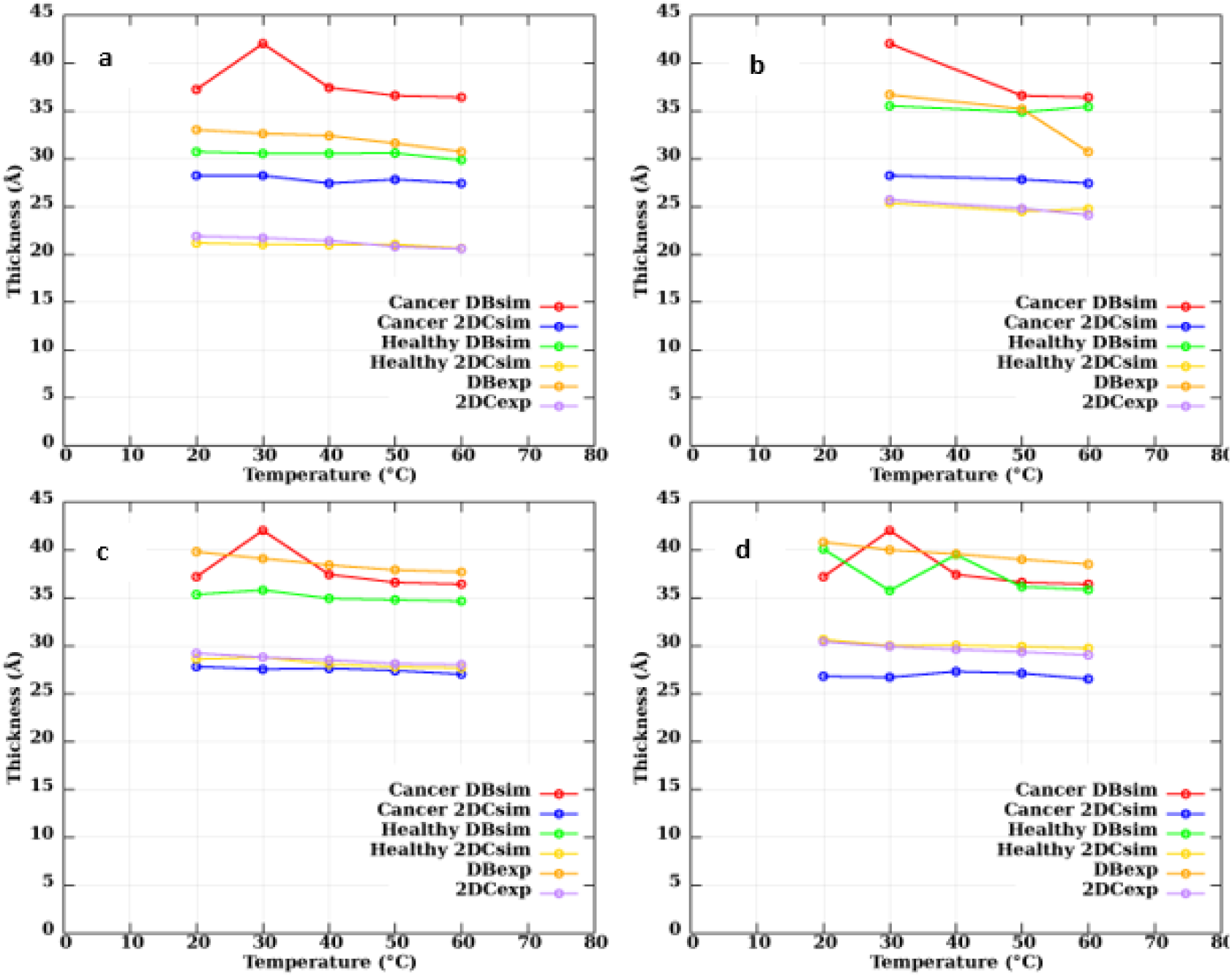
between bilayer thicknesses (DB) and hydrophobic thickness (2DC) for pure and cancer PCs at 20°-60°C to Kučerka et al experimentally-estimated data ^27^. a) Saturated DLPC b) Saturated DMPC c) Unsaturated POPC d) Unsaturated SOPC.

### 3.5 2D Radial Distribution Function

Figure. 7 displays the 2D-Radial Distribution Functions (2D-RDFs) of pure and cancer PCs at temperatures 50°C and 60°C. The 2D-RDF provide valuable insights into the structural properties of PC components, particularly in terms of local order and lipid packing. The 2D-RDFs for both pure and cancer PCs show significant overlap as temperature increases and fluctuates. Fig 7, Fig S.4 and Fig S.5, suggesting that any differences in local packing between the two are not substantial with increasing and varying temperatures. Additionally, at higher temperature of 60°C, a slight increase is observed in the peak of DLPC Fig.5b. A few minor variations in the maximum g(r) values were observed between cancer PCs, other pure PCs and DLPC, with a maximum difference of 0.1 between the two. The peaks for all 2D-RDF functions occurs at approximately 7 Å with a g(r) value of around 1.4, with intermediate positions observed at 8 Å and a well at 10 Å with smaller peaks between 12 Å and 14 Å before plateauing at about 20 Å. Fig S.4 and Fig S.5 show large fluctuations and the presence of noise of the (2D-RDFs) due to small number of lipids in the cancer PC component.

**Fig.7 The.**
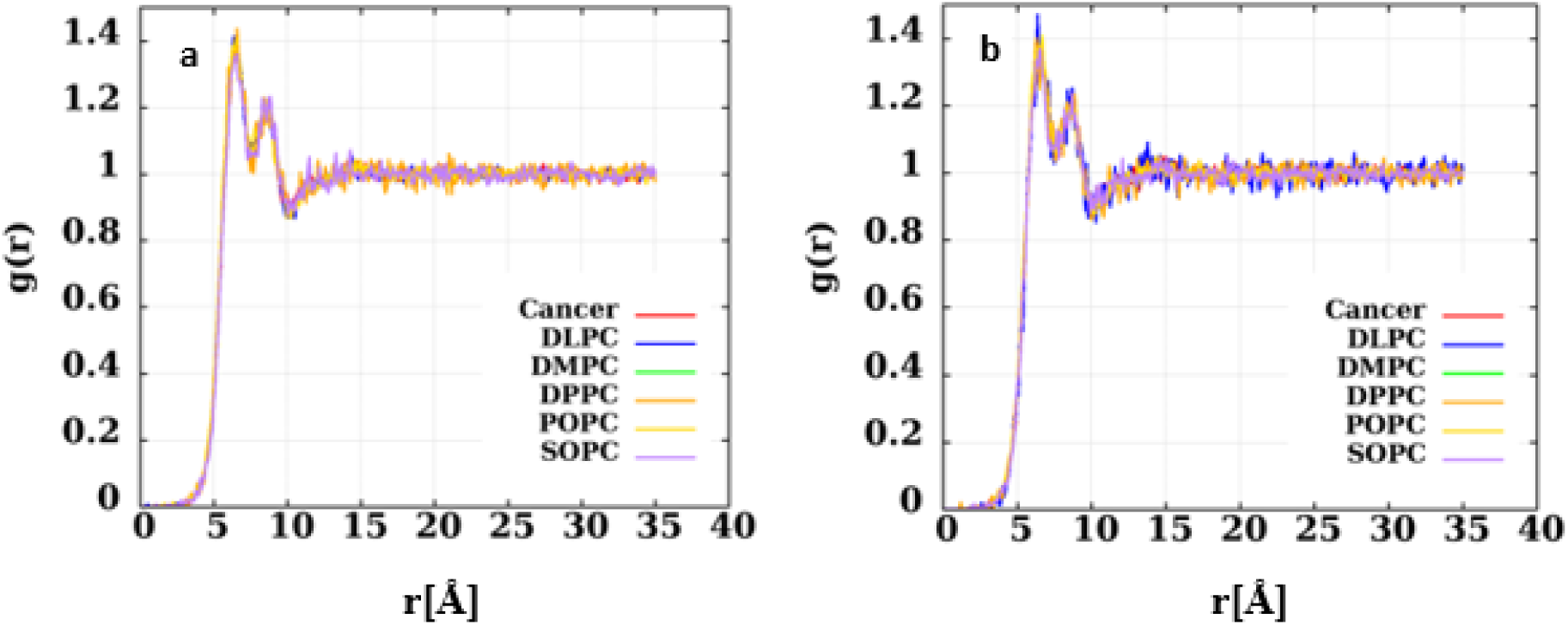
2D radial distribution function a) Mixed Cancer and Pure PC components at 50°C b) Mixed Cancer and Pure PC components at 60°C.

## 4 Summary

Molecular dynamics (MD) calculations were used to analyze splenic pure and cancer PCs of a 5-lipid phosphocholine system, which had compositions determined and derived experimentally. The calculated properties of the PC models are in agreement with experimental pattern and data. Significantly, cancer PCs were found to have a larger surface area per lipid than those of the pure PCs. This indicated that they were bulkier and softer than pure PCs in the lateral direction. In general, cancer PCs had lower SCD values compared to pure PCs. This trend was also seen as temperature increased: the higher the temperature, the less ordered the PCs became. These results indicate that the order parameter of cancer PCs were significantly lower than that of pure PCs. Experimental results for the thickness of the pure PC bilayer (DB) and hydrocarbon region (2DC) are compared with simulation data with evidence of agreement. The results from the electron density studies of the cancer PCs show a decrease in the bilayer thickness as the temperature increases which shows that at higher temperatures, the bilayer thickness decreases for both the pure and cancer PCs. These findings are consistent with each other, supporting this conclusion. The electron density showed less structure for cancer PCs; their first peak was lower and broader while the second and third peaks weren’t clearly visible. The peak of the total EDP for the five-lipid PC system indicates that temperature increase results in thinning of the cancer PC in this lipid system. Looking at all these results together, we hypotize that cancer PCs are more porous and permeable than pure PCs, although this should form a basis for further research with mixed and asymmetric PC bilayers. Our findings demonstrate that the overall structural and dynamical characteristics of the 5-lipid pure phosphocholine bilayer system, obtained through MD simulations, closely match those observed in experimental data. These results validate the significance of using the C36FF model for simulating liquid crystalline bilayers of cancer and pure phosphorlipids with choline only head groups and various chain types at wide ranges of temperature. The slight deviation from experimental data on the surface area per lipid and bilayer thickness observed in the saturated and unsaturated pure PCs at high temperatures could be attributed to the limitations of the C36 force fields and water model used in the simulations. In our future research, we will compare a more comprehensive model of mixed pure mixed with mixed cancer PC phospholipids. This will enable us to observe any potential variance from our current research and expand our analysis to additional phospholipid headgroups.

## Acknowledgment

Financial support was provided by the National Science Foundation and the American Society of Engineering Education. Grant No: EEC 2127509

## 5 Declarations

Conflict of interest: The authors affirm that they have no competing financial interests or personal ties that could have affected the work included in this paper

## SUPPLEMENTARY MATERIAL

### S.0 Calculations

In this study, we assigned lipid molecules with fatty acid tails to specific headgroups based on our understanding of the fatty acid components found in pure and cancer phosphatidylcholine (PC)^1–2^. The sn-1 chain was typically a saturated fatty acid with 16 or 18 carbons, such as palmitic or stearic acid residues, while the sn-2 chain tended to be an unsaturated fatty acid with varying carbons and levels of unsaturation. This unsaturation often took the cis form, causing the chain to bend at that point. It is believed that these cis double bonds are formed through a specific pathway during the biological synthesis of fatty acids. The fatty acid composition and molar percentage ratios of PC were obtained from experimental data^1–2^. Using the method and research conducted by Andoh et al^3^ and the given information above, we were able to determine the number of lipid value and tails for the pure and lymphoma PCs. Although there may be some subjectivity in how we combined different tail chains, our current model closely reflected the data from previous experiments. Based on the experimental data from the normal splenic PC, which has a molar ratio of 0.71 and a phosphatidylcholine composition of 51% of phospholipids, we calculated 0.71 x 51.3 = 36Nip/leaflet for the pure PC. Since this is a simple bilayer model representation, we adopted this for all the 5-lipid composition. For the lymphoma splenic PC, which has a molar ratio of 0.75 and a phosphatidylcholine composition of 58.3%, we calculated 0.75 × 58.3 = 40Nip/leaflet, but modelled it with 70Nip/leaflet to obtain good representation and meaningful result for mixed bilayer simulation.

**Table S.1.**
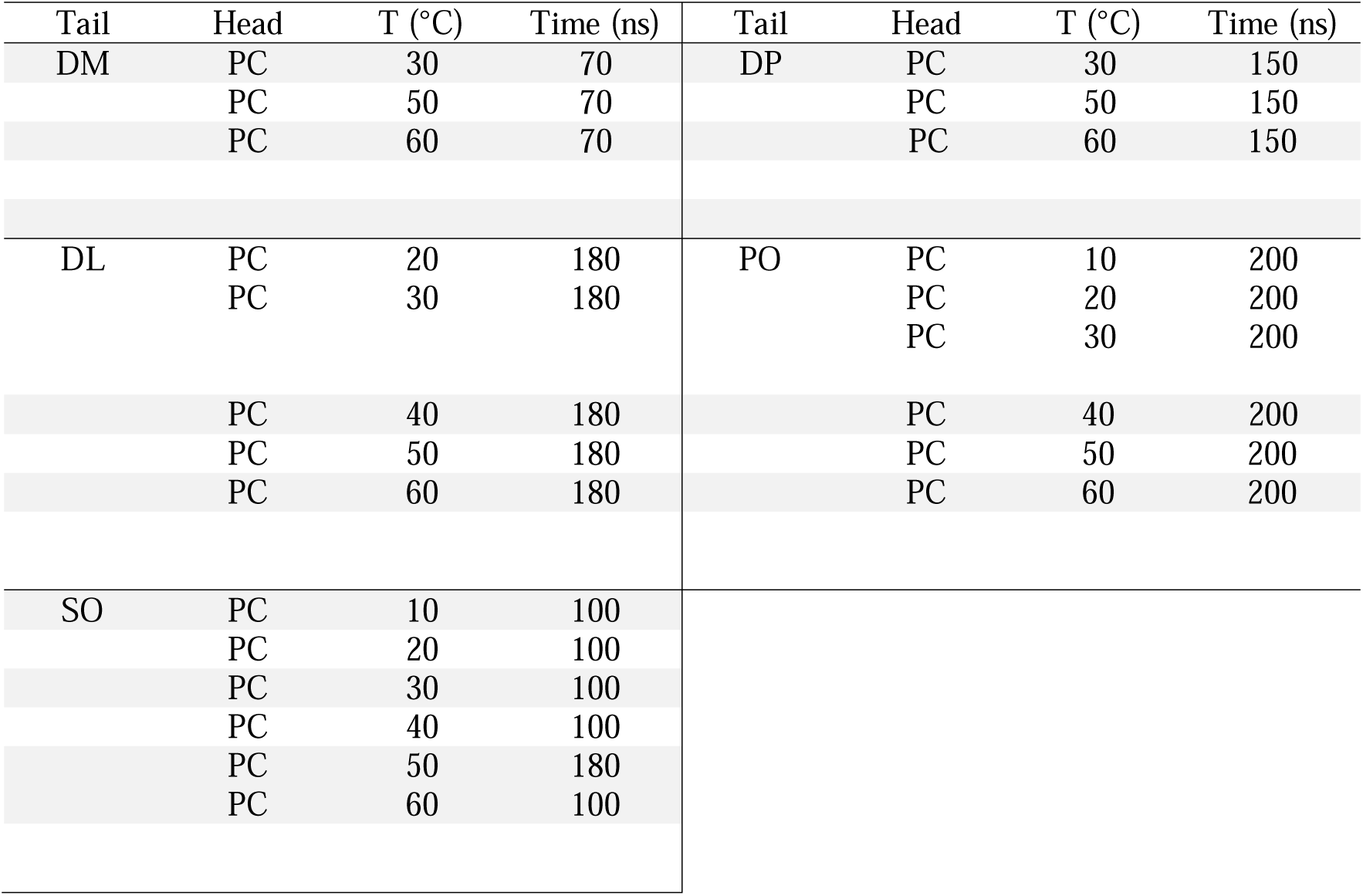
Simulation parameters of pure PC and Leaflet configuration of pure and cancer PC model. Pure PC: (N_Lip/leaflet_) = 36upper:36lower Cancer PC: (N_Lip/leaflet_) = 70upper:70lower.

**Fig S.1.**
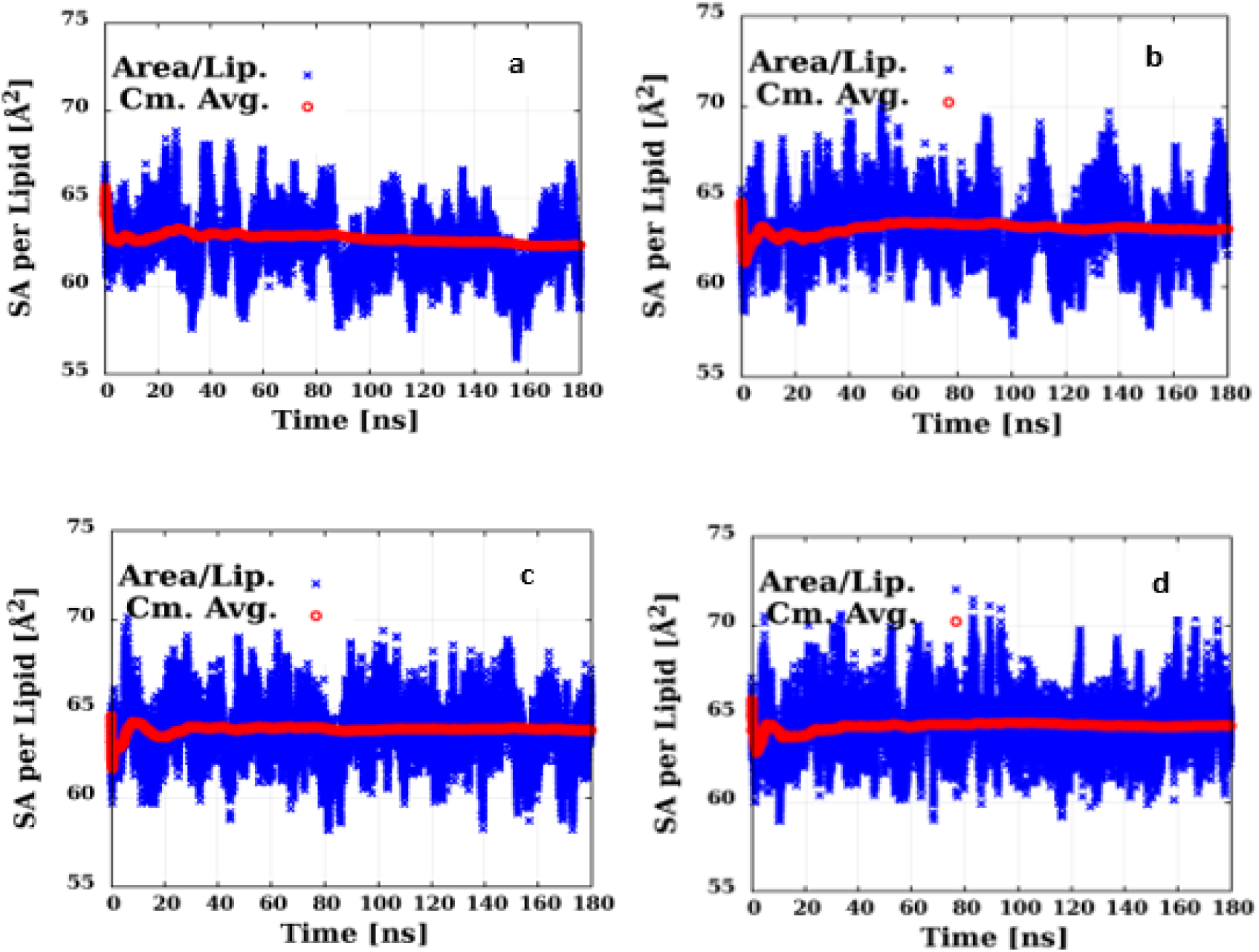
Fig 1. Equilibrated pure model of DLPC SA/lip for a) 30°C b) 40°C, c) 50°C and d) 60°.

**Fig S.2.**
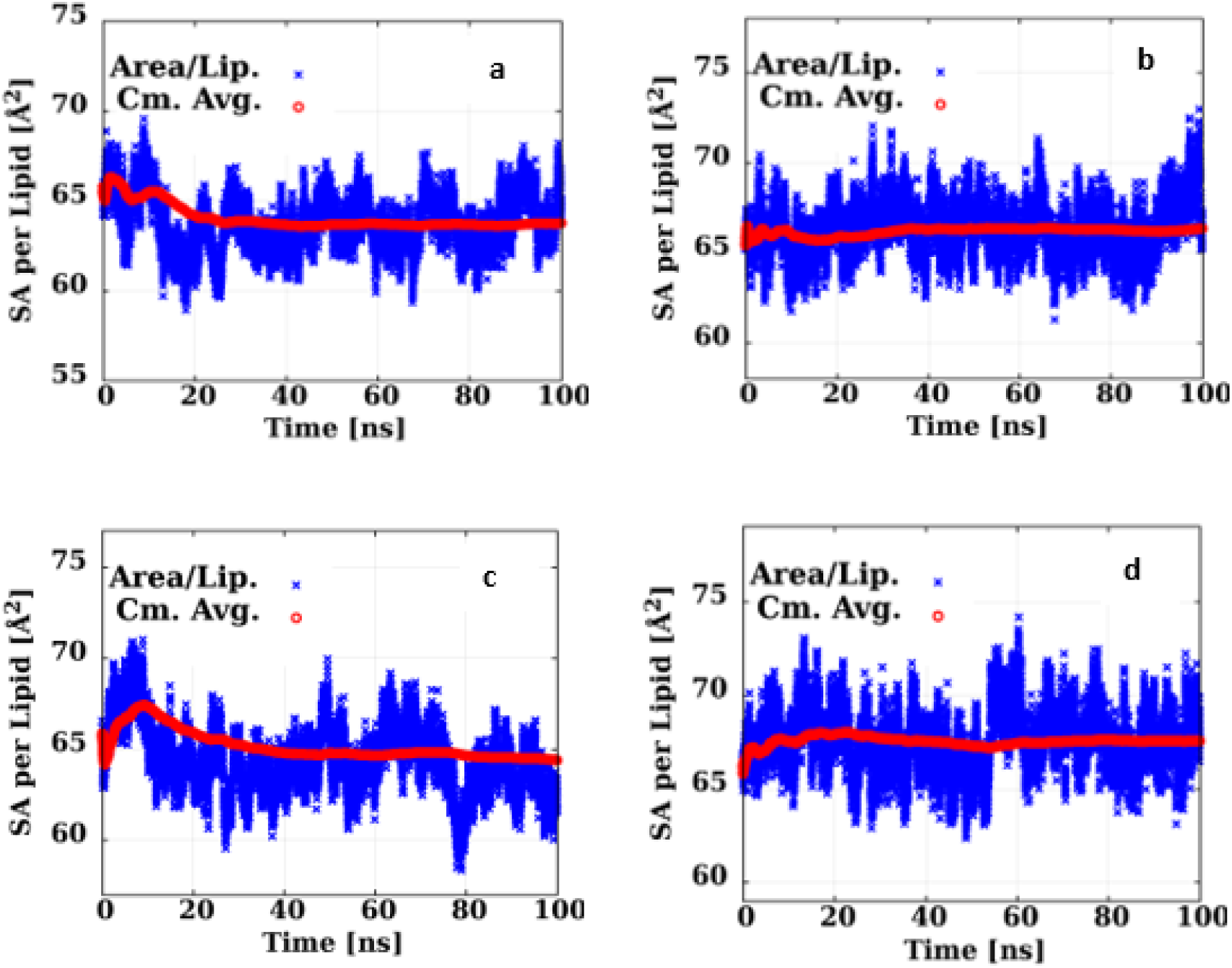
Equilibrated mixed cancer model of DLPC SA/lip for a) 30°C b) 40°C, c) 50°C and d) 60°.

**Table S.2.** SCD values of sn-1 for pure DMPC at 30°-60□ as a function of carbon number.

| c2 | 30°C | 50°C | 60°C |
| --- | --- | --- | --- |
| 2 | 0.0906677 | 0.087551 | 0.0939482 |
| 2 | 0.133427 | 0.108846 | 0.124476 |
| 3 | 0.224353 | 0.199364 | 0.206549 |
| 4 | 0.22727 | 0.203769 | 0.208623 |
| 5 | 0.23927 | 0.212937 | 0.215884 |
| 6 | 0.232493 | 0.204029 | 0.207254 |
| 7 | 0.231274 | 0.201541 | 0.200776 |
| 8 | 0.217098 | 0.188932 | 0.186212 |
| 9 | 0.208971 | 0.178476 | 0.175546 |
| 10 | 0.189364 | 0.161668 | 0.157958 |
| 11 | 0.177903 | 0.147187 | 0.142261 |
| 12 | 0.150818 | 0.121825 | 0.117027 |
| 13 | 0.124297 | 0.097859 | 0.0950921 |
| 14 | 0.0323989 | 0.024098 | 0.0236953 |

**Table S.4.** SCD values of sn-1 for tumor DMPC at 30°-60□ as a function of carbon number.

| **c3** | **30°C** | **50°C** | **60°C** |
| --- | --- | --- | --- |
| **2** | 0.231672 | 0.223057 | 0.224342 |
| **3** | 0.178516 | 0.164537 | 0.168361 |
| **4** | 0.207888 | 0.19241 | 0.194196 |
| **5** | 0.206503 | 0.189654 | 0.187116 |
| **6** | 0.215744 | 0.199089 | 0.192181 |
| **7** | 0.206673 | 0.184499 | 0.17938 |
| **8** | 0.204451 | 0.180885 | 0.176612 |
| **9** | 0.189353 | 0.162217 | 0.158928 |
| **10** | 0.182482 | 0.151504 | 0.149873 |
| **11** | 0.162512 | 0.128246 | 0.125668 |
| **12** | 0.144512 | 0.112466 | 0.109327 |
| **13** | 0.107211 | 0.082205 | 0.078471 |
| **14** | 0.035699 | 0.026264 | 0.02527 |

**Table S.4.** SCD values of sn-2 for pure DMPC at 30°-60□ as a function of carbon number.

| **c2** | **30°C** | **50°C** | **60°C** |
| --- | --- | --- | --- |
| **2** | 0.090879 | 0.098423 | 0.087191 |
| **2** | 0.0851073 | 0.113691 | 0.114684 |
| **3** | 0.192091 | 0.21373 | 0.197568 |
| **4** | 0.208255 | 0.226907 | 0.203927 |
| **5** | 0.222195 | 0.234829 | 0.212565 |
| **6** | 0.219154 | 0.230324 | 0.198197 |
| **7** | 0.212622 | 0.226918 | 0.196936 |
| **8** | 0.1991 | 0.209954 | 0.183078 |
| **9** | 0.185268 | 0.200754 | 0.172375 |
| **10** | 0.160473 | 0.182044 | 0.15919 |
| **11** | 0.147619 | 0.169941 | 0.146105 |
| **12** | 0.122606 | 0.144448 | 0.119366 |
| **13** | 0.0976243 | 0.125479 | 0.0961016 |
| **14** | 0.0243851 | 0.032046 | 0.0269422 |

**Table S.5.** SCD values of sn-2 for tumor DMPC at 30°-60□ as a function of carbon number.

| c3 | 30°C | 50°C | 60°C |
| --- | --- | --- | --- |
| 2 | 0.206068 | 0.232659 | 0.229893 |
| 3 | 0.153662 | 0.188476 | 0.167414 |
| 4 | 0.187752 | 0.216789 | 0.194127 |
| 5 | 0.183869 | 0.205895 | 0.190888 |
| 6 | 0.197354 | 0.214816 | 0.203991 |
| 7 | 0.184085 | 0.203409 | 0.191111 |
| 8 | 0.183658 | 0.20139 | 0.186559 |
| 9 | 0.172757 | 0.174722 | 0.171784 |
| 10 | 0.170063 | 0.165321 | 0.159385 |
| 11 | 0.144827 | 0.1471 | 0.135202 |
| 12 | 0.12292 | 0.131445 | 0.117827 |
| 13 | 0.09509 | 0.093321 | 0.0923958 |
| 14 | 0.035484 | 0.029855 | 0.0317275 |

**Fig.S.2.**
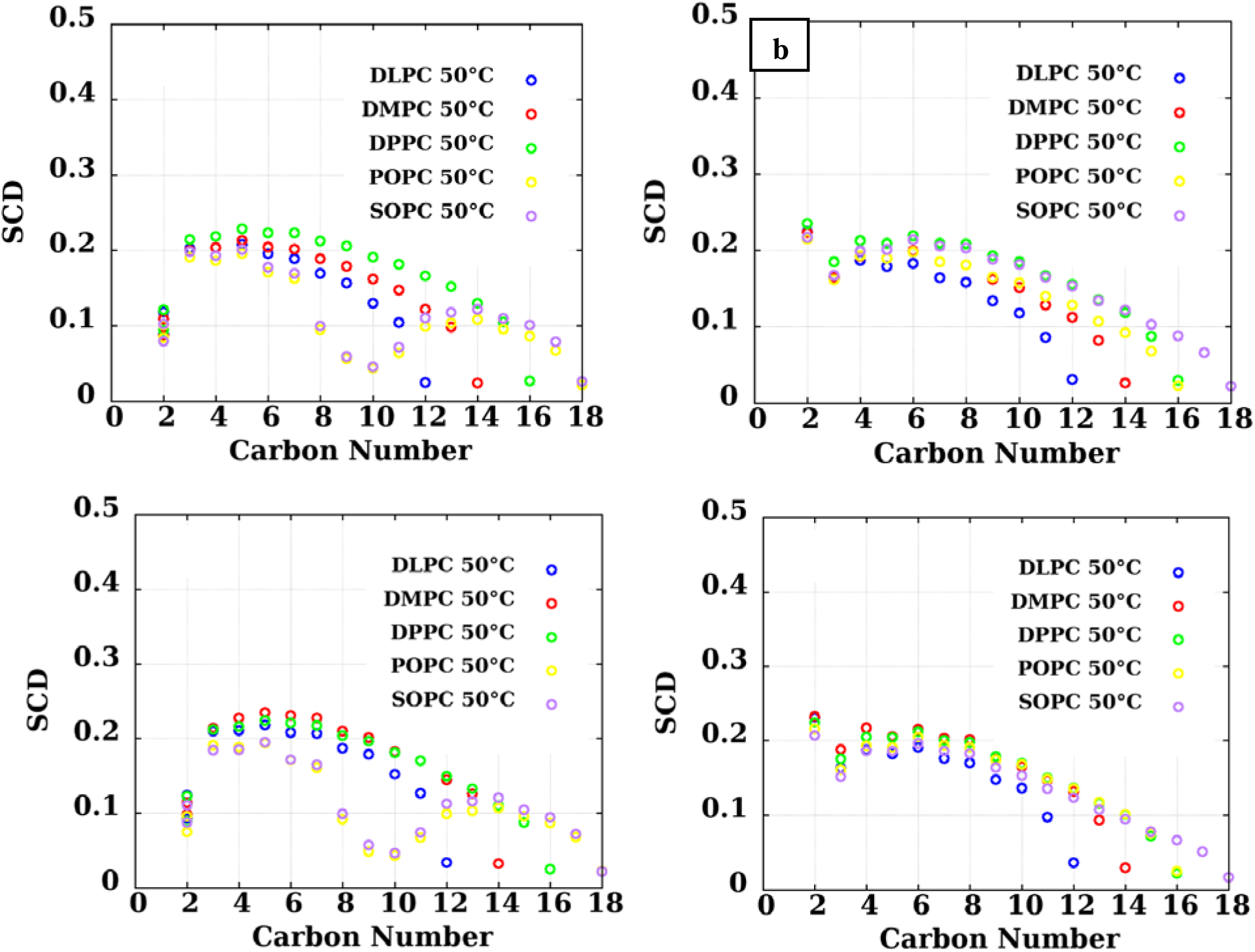
The simulated SCD of five lipid PC system at 50°C (pure DLPC-blue Circle, pure DMPC-red Circle, pure DPPC-Green Circle, pure POPC-Yellow Circle, pure SOPC-Purple Circle. Pure Sn-1 b) Pure Sn-2 c) cancer Sn-1 d) Cancer Sn-2.

**Table S.6.** – Experimental and Simulation bilayer thicknesses consisting of overall bilayer thickness (*D*_B_) and hydrophobic thickness (2*D*_C_) for pure five lipid membrane system.

| Lipid | T(°C) | $DB_{\text{Csim}} (\text{\AA})$ | $DB_{\text{exp}} (\text{\AA})$ | $2D_{\text{Csim}} (\text{\AA})$ | $2D_{\text{Cexp}} (\text{\AA})$ |
| --- | --- | --- | --- | --- | --- |
|  |  | Pure | Pure | Pure | Pure |
| **POPC** | 20°C | 35.35 | 39.8 | 28.6 | 29.2 |
|  | 30°C | 35.79 | 39.1 | 28.76 | 28.8 |
|  | 40°C | 34.91 | 39.1 | 28.04 | 28.5 |
|  | 50°C | 34.8 | 37.9 | 27.75 | 28.1 |
|  | 60°C | 34.65 | 37.7 | 27.64 | 28 |
| **SOPC** | 20°C | 40.08 | 40.8 | 30.61 | 30.4 |
|  | 30°C | 35.74 | 40 | 30.02 | 29.9 |
|  | 40°C | 39.5 | 39.6 | 30.05 | 29.6 |
|  | 50°C | 36.14 | 39 | 29.92 | 29.3 |
|  | 60°C | 35.84 | 38.5 | 29.74 | 29 |
| **DLPC** | 20°C | 30.7 | 21.18 | 33 | 21.9 |
|  | 30°C | 30.51 | 21.04 | 32.6 | 21.7 |
|  | 40°C | 30.52 | 21 | 32.4 | 21.4 |
|  | 50°C | 30.59 | 21.05 | 31.7 | 20.8 |
|  | 60°C | 29.87 | 20.64 | 30.64 | 20.6 |
| **DMPC** | 30°C | 35.48 | 25.36 | 36.7 | 25.7 |
|  | 50°C | 34.88 | 24.46 | 35.2 | 24.8 |
|  | 60°C | 35.41 | 24.7.6 | 30.7 | 24.1 |
| **DPPC** | 50°C | 39.11 | 28.85 | 39 | 28.5 |
|  | 60°C | 38.42 | 28.31 | 38.1 | 27.9 |

**Fig.S.3 Similarities.**
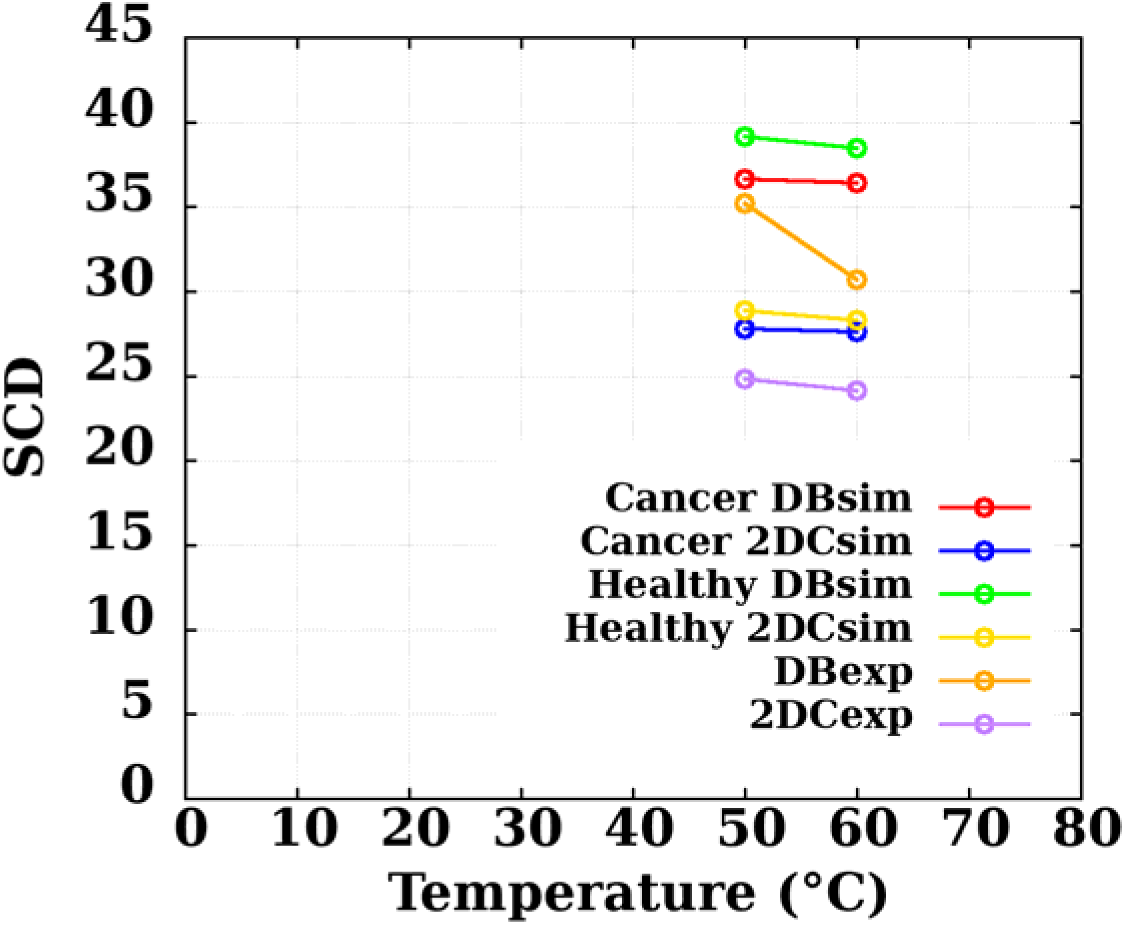
between bilayer thicknesses (DB) and hydrophobic thickness (2DC) for pure and cancer saturated DPPC membrane at 50°C-60°C to Kučerka experimentally-estimated data [4].

**Fig S.4.**
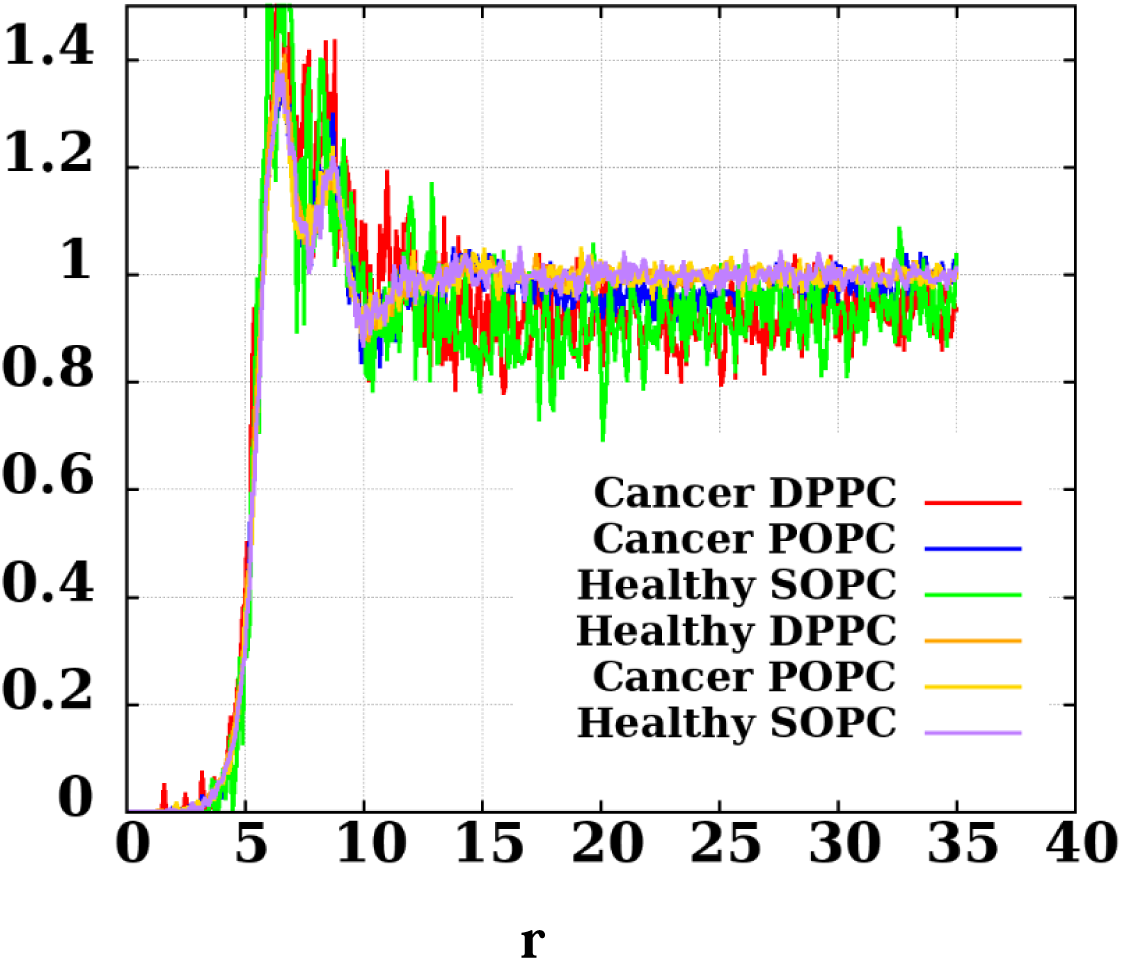
The 2D radial distribution function of Cancer and Pure PCs at various temperatures.

**Fig S.5.**
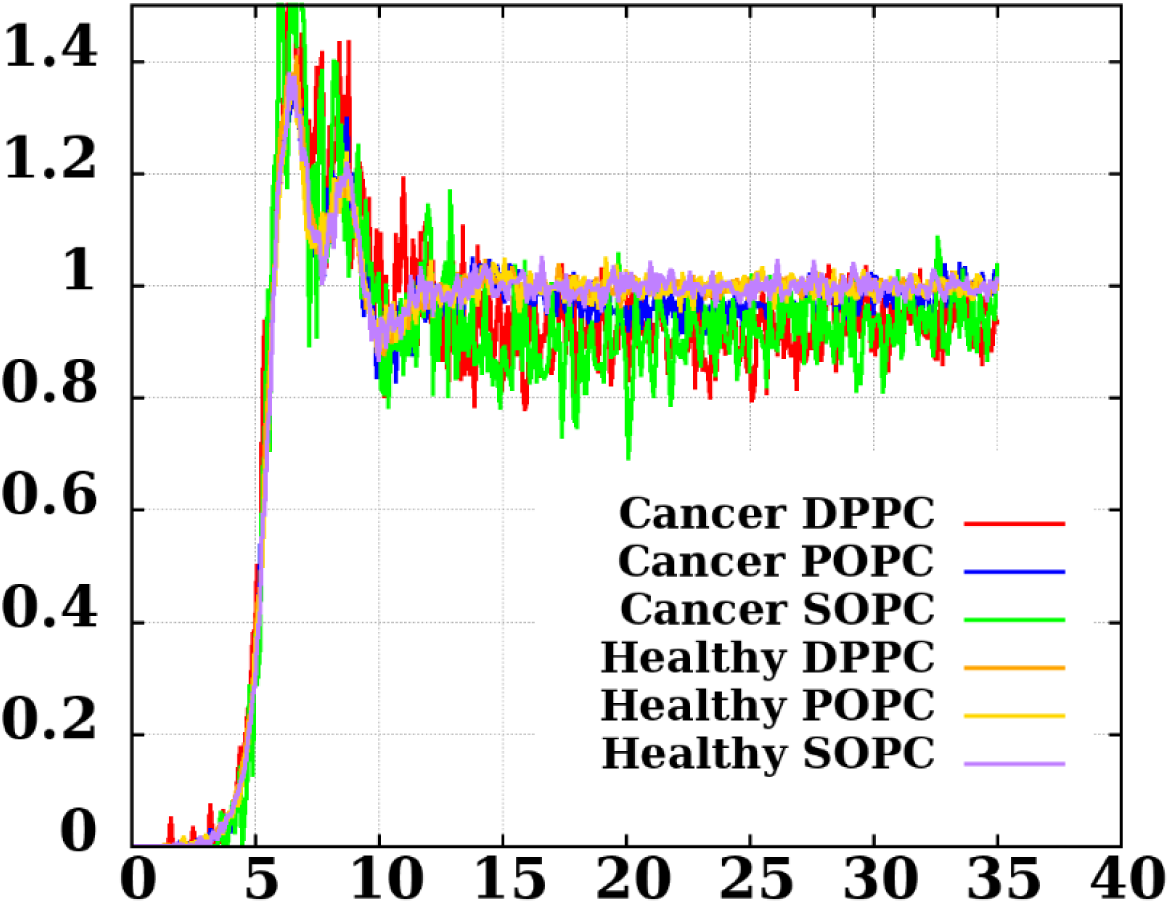
The 2D radial distribution function of Cancer and Pure PCs at various temperatures.

**Fig S.6.**
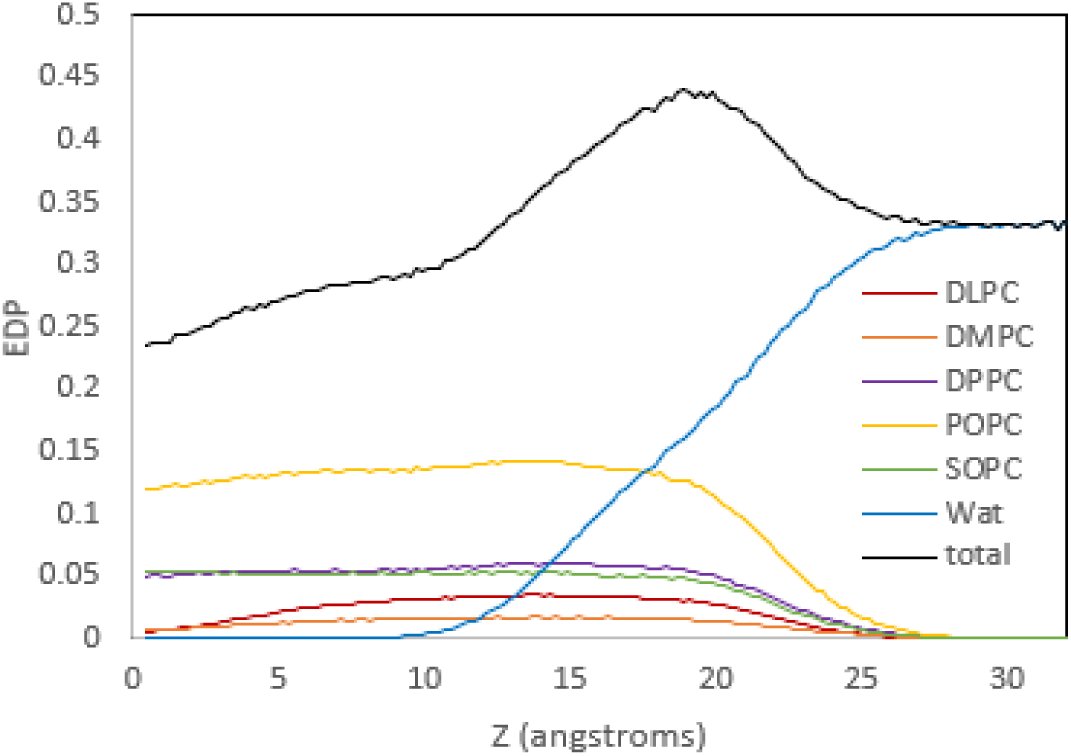
Electron density profiles for 5-lipid cancerous plasma membrane system for temperatures of 50°C.

